# The pro-, but not anti-nociceptive effects of Cannabidiol depend on Trpa1b in larval Zebrafish

**DOI:** 10.1101/2025.10.23.684246

**Authors:** Bryce Lecamp, Quinn Bianucci, Sidhant Rauniyar, Gloria Shen, Kali Esancy, Ben Land, Ajay Dhaka

## Abstract

The promiscuous ligand cannabidiol (CBD) shows promise as an analgesic, but its complex pharmacodynamics has made it difficult to identify its mechanism(s) of action. Numerous putative CBD receptors including cannabinoid receptors CB1 and CB2, as well as Trpv1 and Trpa1—the receptors for capsaicin and mustard-oil (AITC) respectively—have been proposed to contribute to CBD-mediated analgesia. Larval zebrafish have several attributes that lend themselves to inquiries into the biology of nociception. The neural circuits underlying nociception in zebrafish larvae are highly analogous to those found in higher vertebrates. Notably, the small size and optical clarity of zebrafish enable holistic evaluation of analgesic function utilizing behavioral and imaging platforms. Here we report that in larval zebrafish CBD serves both anti- and pro-nociceptive functions. Utilizing place aversion assays as a proxy for nociception, we found that low concentrations of CBD inhibit aversion to noxious chemical stimuli including AITC and acetic acid. Counterintuitively, we found that higher concentrations of CBD potentiated nocifensive behavior as measured by enhanced thermal aversion and increased locomotion. Knockdown of Trpa1b eliminated the algogenic effects of CBD while having no effect on its analgesic properties, as it abolished *trpa1b+* sensory neuron responses to CBD, CBD-evoked thermal hypersensitivity, and increased locomotion in Trpa1b-null animals. Strikingly, CBD profoundly inhibited thermal aversion to noxious heat in Trpa1b-null animals but not in wildtype animals, indicating that CBD-mediated Trpa1b activation can oppose the analgesic properties of CBD. These studies provide a framework to investigate the genetic and neural substrates of CBD-mediated analgesia and nociception.

**Significance statement:** While Cannabis and its component compounds such as delta-9-tetrahydrocannabinol (THC) and cannabidiol (CBD) have long been used to alleviate pain, little is known about their mechanisms of action. Our study provides a framework for investigating the effects of cannabinoids on nociception in a vertebrate, by utilizing behavioral experiments in combination with neuroimaging and in-vitro studies. Together, these approaches allowed us to define the role of one molecular target, Trpa1b, involved in the effects of CBD upon nociception. Critically, activation of this receptor by CBD masks a separate, analgesic effect upon temperature sensation, which suggests a possible explanation for CBD’s inconsistent efficacy across different models of pain.

## Introduction

Pain is the most common reason for patients to seek out medical care in the United States, and chronic pain conditions affect between 10 and 40% of the population (Cohen et al., 2021) Beyond impacting quality of life, pain is debilitating and is proposed to lead to hundreds of billions of dollars of economic loss annually (Smith and Hillner, 2019). The psychological, social, and economic burdens of ineffective pain management are exemplified by the ongoing opioid use epidemic—in part a consequence of our limited options for controlling pain.

Cannabidiol (CBD) is a naturally occurring compound produced by *Cannabis sativa*. Preparations of this plant, which include CBD and other terpenoids—collectively called phytocannabinoids—have been used medicinally and recreationally for millenia (Russo, 2007). Despite anecdotal evidence for its use in alleviating chronic pain, as well as for improving sleep and controlling anxiety, CBD has failed to consistently demonstrate efficacy in clinical trials designed to assess its analgesic potency (Hunter et al., 2018; Vučković et al., 2018; Nitecka-Buchta et al., 2019; van de Donk et al., 2019; Eskander, Md, Mba et al., 2020; Mlost et al., 2020; Urits et al., 2020; Xu et al., 2020; Bergmans et al., 2024; Mohammed et al., 2024; Wang et al., 2025).

Contributing to this confusion, CBD is notorious even among phytocannabinoids for its complex pharmacology. While it has generally been shown to have low activity at the canonical cannabinoid receptors, CB1 & CB2, CBD interacts with metabotropic and ionotropic targets expressed in neurons, and many other cell types (Stella, 2023). Among those implicated in the pain pathway are the TRP channels Trpa1 and Trpv1, both well-established mediators of noxious sensation (Laing and Dhaka, 2016) and activated by CBD in mammals (Bisogno et al., 2001; Jordt et al., 2004).

Zebrafish larvae, *Danio rerio*, have gained popularity as a model organism for biomedical research due to being vertebrates, while sharing the small size, genetic toolkit, and capacities for *in-vivo* imaging found in non-vertebrate model systems such as *Drosophila melanogaster* and *Caenorhabditis elegans* (Costa et al., 2022). Our group and others have demonstrated the conservation in larval zebrafish of critical anatomic, circuit, and molecular mediators of nociception found in mammals, including the function of TRP channels for aversion to noxious chemicals and heat (Prober et al., 2008; Gau et al., 2013; Sneddon, 2019). Using previously validated assays (Curtright et al., 2015; Esancy et al., 2023) for sensitivity to painful stimuli, we sought to determine CBD’s effectiveness as an analgesic in larval zebrafish, as well as potential mechanisms by which it exerts any of its effects.

We demonstrate that at relatively high concentrations CBD can directly activate Trpa1b, one of two zebrafish genes paralogous to mammalian TRPA1, which is expressed in nociceptors innervating the skin. By activating Trpa1b, CBD induces thermal hypersensitivity as well as hyperlocomotion. These nocifensive behaviors are eliminated by genetic and pharmacological knockdown of Trpa1b, but are not affected by loss of Trpv1. Conversely, we show that at relatively low concentrations, CBD reduces aversion to the chemical irritants allyl-isothiocyanate (AITC) and acetic acid. In contrast to CBD’s pro-nociceptive effects at higher doses, reduced aversion to chemical algogens does not require Trpa1b, Trpv1, or the cannabinoid receptors Cnr1 and Cnr2. Further, only fish lacking Trpa1b lose their aversion to the noxious temperature 37.5°C, in the presence of CBD, suggesting that CBD’s effects on Trpa1b masks an antinociceptive effect on temperature sensation.

Collectively, we demonstrate both pro- and anti-nociceptive effects of CBD that depend on concentration and the way in which pain sensitivity is measured. We also show that the pronociceptive properties of CBD can antagonize its antinociceptive effects. We conclude that CBD’s diverse pharmacological activities may functionally oppose each other, and that understanding its analgesic potency requires considering its impact on multiple, potentially counteractive, targets.

## Methods

### Zebrafish husbandry

Adult zebrafish (Danio rerio) were raised with constant filtration, temperature control (28.5 ± 2 °C), illumination (14 h:10 h light–dark cycle, lights on at 9:00 AM), and feeding. All animals were maintained in these standard conditions, and the Institutional Animal Care and Use Committee approved all experiments. Adult zebrafish not used in behavioral experiments were bred in spawning traps (Thoren Caging Systems, Hazelton, PA) from which embryos were collected. Larval zebrafish were raised in petri dishes (Fisher Scientific, Hampton, NH) of E2 medium (EM) with no more than 50 embryos per dish at 28.5 ± 1 °C in an incubator (Fisher Scientific). Embryos were staged essentially as described (<u>Kimmel et al., 1995</u>) and kept until 6dpf. Cannabinoid receptor knockout animals were a gift from Wolfram Goessling (Harvard Medical School) and maintained on the Tubingen background.

### Chemicals

CBD and CBC used in this study was purchased from Cayman Chemical Co Inc. (product # 90080, 26252). TRPA1 antagonist HC-030031 was purchased from Sigma-Aldricht (product # H4415). The following were purchased from Fisher Scientific Co LLC: Allyl isothiocyanate (mustard oil, AC102951000), Ethanol (04-355-226) and Dimethyl Sulfoxide (DMSO, D128-4). All other reagent sources are noted in their respective sections.

### Behavioral Assays

#### Temperature aversion assay

Temperature aversion was performed as previously described (Curtright et al., 2015). Briefly, randomly selected P5/6 larvae were pipetted in 100µL of EM into individual wells of a custom printed 32-well plate. 100µL of 2X solution was added, and then the plate was placed into a 28.5°C incubator for the indicated incubation period. Subsequently, the plate was adhered to the surface of a two-sided hot plate with dH2O. For experiments involving knockout animals, their genotype was assessed after the experiment by HRMA (see below). Room temperature was maintained at 27-28°C.

#### Chemical aversion assay

Chemical place aversion was performed as previously described (Esancy et al., 2023). Briefly, randomly selected P5/6 larvae were pipetted in 100µL of EM into a petri dish, and EM was added to a final volume. An equal volume of 2X Solution was added and the fish were moved to a 28.5°C incubator for the indicated incubation period.

Subsequently, the petri dish and its contents were poured into a square petri dish lined with 0.8% agarose EM on three sides, and 100mM AITC infused into 0.8% agarose EM on a fourth side. Experiments involving knockouts used fish from a cross of homozygous parents. Acetic acid aversion was performed identically, with glacial acetic acid (1% final concentration) in place of AITC. Room temperature was maintained at 27-28°C.

Analysis was performed using FastTrack (Gallois and Candelier, 2021). Briefly, a blank background was created in ImageJ by performing a max projection on the video of an experiment. This background was used in FastTrack, along with custom settings for each video. Data from Fastrack was uploaded into Excel and analyzed using a custom script.

#### Locomotor assay

Locomotion was assessed by pipetting P5/6 larvae into the wells of a 96-well plate mesh-bottom plate, and subsequently moving this plate from embryo media into a bath of the indicated solution kept at room temperature unless otherwise indicated. For experiments involving knockout animals, their genotype was assessed after the experiment by HRMA. Analysis was performed in EthoVision, and data were exported in 10-second bins. Room temperature was maintained at 27-28°C.

#### High Resolution Melt Analysis (HRMA)

HRMA was performed on a BioRad CFX connect with precision melt analysis software. PCR primers (ordered from IDT) flanking the site of an indel were chose to create a 50-150bp amplicon, which was amplified in 10µL with the following settings: 40 cycles, 95c for 15s, 60c for 45s, 72c for 30s. Melt curve images were taken from 70 to 85c in 0.1c increments, 10 seconds per step. The template DNA for each reaction was extracted using the Hot Base method (Meeker et al., 2007), using 1µL per well.

For genotyping of the *trpa1b* Δ7 knockout line, the following primers were used:

Trpa1b HRMA FWD primer: 5’ACTTTGATTTGTGTCAGATATGGG’3
Trpa1b HRMA REV primer: 5’TTAGAGTCTGTGAGGGTCTCC’3

For genotyping of the *cnr1* and *cnr2* knockout lines, the following primers were used:

*cnr1* HRMA FWD primer: 5’GCGGAAACCACCTTCAGA’3
*cnr1* HRMA REV primer: 5’CGTCGTAGCCGATGTCATT’3

*cnr2* HRMA FWD primer: 5’GTGTGGTTTGTGATTCCAAATTC’3
*cnr2* HRMA REV primer: 5’AACAGTACCTGTGGTTGAAATG’3

### Generation of *trpv1* mutant

The *trpv1* nonsense mutant was generated as described previously (Esancy et al., 2018). Briefly, freshly fertilized embryos were microinjected with 100 ng/μL of gRNA and 400ng/μL of Cas9-NLS protein (PNA Bio). A gRNA targeting the sequence 5’AGGCGGTGGATGTCAGGGAGAGG’3 was chosen using CRISPRscan (Moreno-Mateos et al., 2015) to target an early exon with high efficiency and low off-target cutting. This gRNA produced in-vitro with a T7 transcription kit (Invitrogen). Injected animals were grown to adulthood and outcrossed to isolate founders with HRMA (primer sequences below). The mutation was verified using Sanger sequencing (GENEWIZ from Azenta Life Sciences), and a single founder was outcrossed to generate the knockout line. This line has a 16 base pair deletion beginning at R239, causing an early stop after only 246 amino acids and truncating the predicted protein product before any transmembrane helices (Liao et al., 2013).

Trpv1 hrma fwd primer: 5’ CAGCAAGACATTGTGGACTTTC’3
Trpv1 hrma rev primer: 5’ TTTCAGGGCTATTATCAGCTACG’3
Trpv1 sequencing fwd primer: 5’ CCAACTATGACGACTTCCACTAC’3
Trpv1 gRNA 1 in-vitro transcription primer:
5’taatacgactcactataGGGCGGTGGATGTCAGGGAGgttttagagctagaa’3

To confirm the loss of mRNA associated with the mutation and presumed nonsense-mediated decay (El-Brolosy et al., 2019), the *trpv1* mRNA was visualized using colorimetric in-situ hybridization with a whole-cDNA antisense probe, as described previously (Gau et al., 2013). Briefly, offspring from a cross of parents heterozygous for the mutation were exposed to 0.003% PTU from 24hrs post fertilization onwards. At 3dpf, larvae were euthanized and DNA was extracted from a small section of the tail for genotyping. WT and KO larvae were then fixed in formalin prior to in situ hybridization. For imaging, stained fish were mounted in agarose and imaged with brightfield settings on a Keyence BZ-X800 Light Microscope (Keyence), image stacks were processed with Helicon Focus (Helicon Soft LTD).

### In vivo Ca2+ imaging of trpa1b+ neurons

To drive expression of reporter genes in trpa1b-expressing cells, we amplified roughly 5kb of sequence immediately upstream of the trpa1b start codon using PCR (primers below). This sequence was cloned upstream of the zebrafish beta-actin minimal promoter, GCaMP6s, and a sv40 poly-adenylation sequence, all flanked by iTol2 arms in the plasmid pDest. Injection of this plasmid revealed specific expression of GCaMP6s in the Trigeminal ganglia and Rohon-Beard neurons. Calcium imaging confirmed that all neurons were AITC-responsive.

trpa1b promoter forward: 5’tacctatcatatagatcttt’3
trpa1b promoter reverse: 5’gtcactcagtatgaagctgg’3

AB or *trpa1b* Δ7 knockout embryos were injected with pTol2-*trpa1b:GCaMP6s* (15 ng/μl) and Tol2 transposase (50 ng/μl) to image activity in *trpa1b*+ neurons. At 72 hpf, injected larva were screened for green fluorescence in the trigeminal ganglia under Tricaine anesthesia. At 120 or 144 hfp in ice-cold EM and immobilized by a harp (Warner Instruments) larvae were paralyzed by injecting α-bungarotoxin protein (Sigma) into the chest cavity using microinjection needles pulled on a Flaming-Brown Micropipette Puller (model P-87, Sutter Instrument Co., Novato, CA) and a Picrosprizter II microinjection apparatus (General Valve Corporation, Fairfield, NJ). After injection, these larvae were kept at room temperature in a petri dish before imaging, which occurred within 5 hours of paralysis.

After paralysis, larvae were mounted in 3% low melt agarose prepared in EM onto coverslips, which were then placed into a perfusion chamber (Warner Instruments). The agarose immediately surrounding the head was cut away with a scalpel to ensure maximal exposure to chemical stimuli.

The perfusion chamber was placed onto the stage of an Olympus IX81 inverted fluorescence microscope equipped with real-time acquisition and graphing software (MetaFluor; Molecular Devices). Larvae were imaged at 10x with a GFP filter, each frame was taken under a 300 ms exposure at 0.25 frames/s. Embryo medium containing 1% DMSO and 0.2% EtOH by volume was constantly perfused from an in-line perfusion system. The temperature was adjusted with an in-line heater/cooler (Warner Instruments) and maintained between 23 and 25°C.

The resulting images were uploaded into FIJI, cropped to include only the cells of interest, and registered to the first frame using TurboReg. ROIs were drawn over GcaMP6s+ and their average intensity value was extracted and exported into excel. After background subtraction, changes in fluorescence intensity over time were calculated relative to the average prestimulus baseline as ΔF/F.

For quantification of average responses, we set a threshold for activation at a 30% increase over baseline (0.3 ΔF/F). We summed the total area between the ΔF/F value and the threshold (0.3) for every frame over the course of a treatment block, then divided this value by the total time of the block, giving an ‘average response’ for a given treatment, which is reported as an arbitrary unit (AU). Responder neurons were quantified as having a greater response in AU than 0.002.

### Ratiometric calcium imaging

Human embryonic kidney (HEK) 293T cells were cultured in DMEM (Invitrogen) supplemented with fetal bovine serum and antibiotics (penicillin/streptomycin), and passaged every 2–3 days.

Calcium imaging was performed essentially as described previously (Dhaka et al., 2009). HEK cells were transiently transfected with Trpa1b.pcDNA3.1 and pIRES-eGFP or pIRES-eGFP alone. Transfections were incubated overnight at 37°C and then transferred to incubate at 30°C approximately 18 hours prior to experiment. The buffer solution for all experiments was 1× Hanks’ balanced salt solution (HBSS; Invitrogen) and 10 mm HEPES (Invitrogen). All data presented was captured exclusively from cells expressing eGFP. Cells not expressing eGFP were also recorded but data was not shown as they did not differ significantly from eGFP-expressing cells.

Averaged traces represent mean ± s.e.m. All reported fluorescence values of each cell were normalized to the fluorescence of that cell during the initial baseline wash period.

### Statistical Analysis

Statistical analyses were performed using GraphPad Prism 10. Statistical significance was calculated using student’s t-test for comparisons between two groups after testing for normality, or a Mann-Whitney test. For comparisons between three groups or more, one or two-way ANOVA with appropriate Tukey or Dunnet corrections for multiple comparisons were used. For linear regression, the simple linear regression function was used with a test for significance of the slope. P values less than 0.05 were considered statistically significant. The data are reported as the mean ± standard error of the mean.

## RESULTS

### CBD provokes *trpa1b*-dependent thermal hyperalgesia and hyperlocomotion in larval zebrafish

The Trpa1 agonist allyl-isothiocyanate (AITC) is commonly used in models of sensitized pain, where activation of Trpa1 can provoke hypersensitivity across pain modalities (thermal, mechanical, etc) (Bautista et al., 2006). AITC induces sensitization to heat in larval zebrafish in a Trpa1 dependent manner; a temperature which is only mildly aversive at baseline, 31.5°C, becomes strongly aversive after treatment with AITC, causing animals to spend a significantly greater proportion of time at their rearing temperature, 28.5°C (Curtright et al., 2015). This sensitization to heat is alleviated by pre-treatment with drugs known to be analgesic, such as the opioid buprenorphine, as well as by the Trpa1 antagonist HC-030031 (Curtright et al., 2015).

We utilized this model of sensitized nociception to determine whether CBD acts as an analgesic in larval zebrafish (Fig. 1A). We incubated larval zebrafish with vehicle (1% DMSO + 0.2% ethanol in embryo media, EM) or CBD for 40 minutes, adding a sensitizing dose of AITC 10 minutes before assessing thermal sensitivity by setting one half of an oval arena to 28.5°C and the other half to a test temperature, in this case 31.5°C. After treatment with vehicle, larvae spent an average of 53.1% ± 3.77 of their time on the 28.5°C side of the arena, indicating no aversion to 31.5°C. When given a sensitizing dose of AITC in addition to the vehicle treatment, the animals spent instead 82.3% ± 2.24 of their time at 28.5°C—avoiding 31.5°C significantly. Rather than reducing the sensitization induced by AITC, treatment with 4µM CBD only exacerbated its effect (Fig. 1B), bringing aversion to 92.9% ± 1.62, while lower doses had no effect at all. Further, at a slightly higher dose (5µM), adding CBD simultaneously with AITC and incubating for 10 minutes was sufficient to induce similar sensitization to 4µM for 40 minutes (Fig. 1D), indicating that CBD’s effects on this assay do not require prolonged exposure.

**Figure 1).**
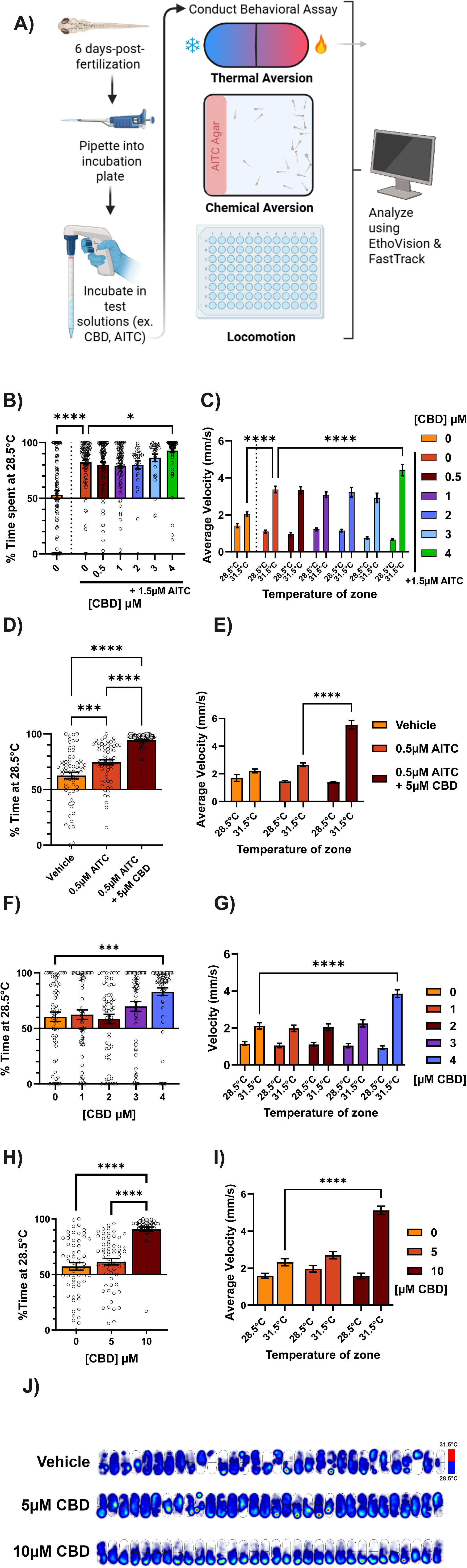
CBD induces thermal hypersensitivity. a) Graphical abstract for behavioral experiments. P5/6 larvae are pipetted into experimental apparati in embryo media, to which is added the test solution. After an incubation period, animals’ behavior is monitored by video and analysis is performed through an automated pipeline. b) Dose response for 40 minute incubation of CBD + 10 minute incubation with 1.5µM AITC prior to testing temperature preference for 28.5°C vs 31.5°C. Data represent average time spent on the 28.5°C side across all fish as a percentage of total trial duration (4 minutes). AITC induced thermal hypersensitivity, and CBD is unable to rescue this effect. At 4µM, CBD increased thermal hypersensitivity above that induced by AITC. Left 0µM CBD treatment is vehicle only, right 0µM over the bar is vehicle plus AITC. N = 64 larvae per group. c) Average velocity (mm/s) of larvae in either temperature zone for experiment in panel B. AITC induced hyperlocomotion specifically on the 31.5°C side and 4µM CBD, but not lower concentrations, significantly increased this effect. d) Effect of 10 minute incubation of CBD + 0.5µM AITC prior to testing temperature preference for 28.5°C vs 31.5°C. 5µM CBD causes increased thermal hypersensitivity above AITC alone. N = 64 larvae per group. e) Average velocity (mm/s) of larvae in either temperature zone for experiment in panel D. 10 minute treatment with 5µM CBD + AITC, but not AITC alone, causes hyperlocomotion specifically on the 31.5°C side. f) Dose response for 40 minute incubation of CBD alone prior to testing temperature preference for 28.5°C vs 31.5°C. Data represent average time spent on the 28.5°C side across all fish as a percentage of total trial duration (4 minutes). 4µM CBD itself induces thermal hypersensitivity to 31.5°C. N = 64 larvae per group. g) Average velocity (mm/s) of larvae in either temperature zone for experiment in panel F. 4µM CBD induced hyperlocomotion specifically on the 31.5°C side, with lower doses having no effect. h) Dose response for 10 minute incubation of CBD alone prior to testing temperature preference for 28.5°C vs 31.5°C (n = 64 per group) as a percentage of total time (4 minutes). 5µM CBD alone has no effect on aversion to 31.5°C, but 10 µM CBD induces profound thermal hypersensitivity. N = 64 larvae per group. i) Average velocity (mm/s) of larvae in either temperature zone for experiment in panel H. 10, but not 5µM CBD alone induced hyperlocomotion specifically on the 31.5°C side. j) Heat maps indicating the cumulative position of the animals over the course of the experiment in panel G. Brighter/hotter colors indicate more time spent in those pixels. The 28.5°C zone is below the dark line across the center of the arena, and the 31.5°C zone is above this line. Vehicle and 5µM CBD treated animals explored both sides of the arena, while 10µM treated animals generally avoided the 31.5°C side. *p < 0.05; **p < 0.01; ***p < 0.001; ns, not significant. Error bars represent SEM. Panels B, D, F, H used one-way ANOVA; C, E, G, I, used two-way ANOVA.

Larvae have a higher average velocity on the side of the arena they avoid, characteristic of escape responses after encountering noxious stimuli (Esancy et al., 2023). While vehicle-treated animals displayed only a 45% increase in velocity (1.42 ± 0.114 to 2.06 ± 0.134 mm/s) from the 28.5 to the 31.5°C side, AITC treatment tripled the average velocity on the 31.5°C side (1.10 ± 0.071 to 3.37 ± 0.174 mm/s) after 40 minutes of vehicle (Fig. 1C). CBD treatment exacerbated this effect in a dose-dependent manner, causing an even greater increases in average velocity on the 31.5°C side in the presence of 4µM CBD (Fig. 1C,E). Therefore, like its effect on aversion to 31.5°C, CBD also potentiates escape responses stimulated by 31.5°C.

Given that CBD is known to activate mammalian TRP channels (Bisogno et al., 2001; Jordt et al., 2004) and could potentiate the effects of AITC on thermal sensitization in zebrafish, we tested whether CBD *alone* could provoke thermal hyperalgesia. When the AITC was removed, 4µM CBD indeed caused hypersensitivity to 31.5°C (60.5% ± 4.28 to 83.1% ± 3.51 time on 28.5°C for 40 minutes of 0 or 4µM CBD, respectively, Fig. 1F,G), while lower doses had no effect on aversion or velocity in the 31.5°C zone. Furthermore, a higher dose (10µM) also caused aversion and increased velocity in the 31.5°C zone when tested after only 10 minutes of pre-treatment (Fig. 1H,I). Indeed, a heatmap of the cumulative position of each fish reveals that 10, but not 5µM CBD incubation for 10 minutes causes a dramatic shift in occupancy towards the 28.5°C side of the arena (Fig. 1J). These results are consistent with studies demonstrating an effect of time on CBD uptake in laval zebrafish, where the 40 minute timepoint is closer to when CBD absorption should be at its peak (Achenbach et al., 2018) while after only 10 minutes CBD concentration is still increasing.

Given that CBD could effectively mimic AITC in the 31.5°C place preference assay, we determined whether Trpa1 was required for CBD’s effects on temperature preference. Co-incubation of CBD with a Trpa1 antagonist, HC-030031, blocked (81.9% ± 2.68 to 54.0% ± 5.58 time spent at 28.5°C) 10µM CBD-evoked thermal hyperalgesia and increased velocity at 31.5°C (Fig. 2A, B). Further, CBD failed to induce thermal hyperalgesia in *trpa1b* homozygous knockout (-/-) animals after both 10 (Fig. 2C, D) and 40 (Fig. 2E, F) minute incubations. Heatmaps for *trpa1b* −/− animals treated for 10 minutes with 10µM CBD show roughly equal occupancy of the 28.5 and 31.5°C zones in contrast to wild-type animals (Fig. 2G).

**Figure 2).**
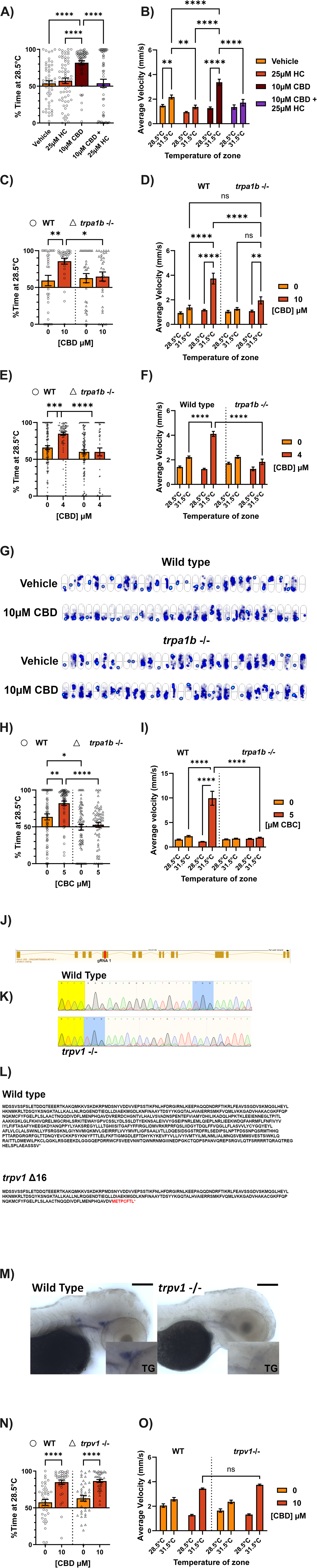
CBD-thermal hypersensitivity is dependent on Trpa1b, but not Trpv1. a) Effect of 10 minute co-incubation of 10µM CBD with the Trpa1 antagonist HC-030031 (25µM) prior to testing temperature preference for 28.5°C vs 31.5°C. HC-030031 does not affect temperature preference on its own, but blocks CBD-induced thermal hypersensitivity. N = 53-62 larvae per group. b) Average velocity (mm/s) of larvae in either temperature zone for experiment in panel A. 10 minute incubation with 10µM CBD induced hyperlocomotion specifically in the 31.5°C zone, which is blocked by co-incubation of with 25μM HC-030031. HC-030031 on its own reduced velocity on the 31.5°C side significantly. c) Effect of *trpa1b* homozygous knockout (triangles) on temperature preference for 28.5°C vs 31.5°C in the presence of CBD. Aversion to 31.5°C stimulated by 10 minute incubation with 10µM CBD is eliminated by knockout of *trpa1b*. N = 32 larvae per group. d) Average velocity (mm/s) of larvae in either temperature zone for experiment in panel C. CBD’s specific increase in velocity on the 31.5 side is eliminated with knockout of *trpa1b*. e) Heat maps indicating the cumulative position of the animals over the course of the experiment in panel C. Brighter/hotter colors indicate more time spent in those pixels. The 28.5°C zone is below the dark line across the center of the arena, and the 31.5°C zone is above this line. Wild-type animals exhibit a large change in preference between vehicle and 10µM CBD treatment, while *trpa1b* knockout animals do not. f) Effect of *trpa1b* homozygous knockout (triangles) on temperature preference for 28.5°C vs 31.5°C in the presence of CBD. Aversion to 31.5°C stimulated by 40 minute incubation with 4µM CBD is eliminated by knockout of *trpa1b*. N = 64-127 larvae per group. g) Average velocity (mm/s) of larvae in either temperature zone for experiment in panel F. CBD’s specific increase in velocity on the 31.5 side is eliminated with knockout of *trpa1b*. h) Effect of *trpa1b* homozygous knockout (triangles) on temperature preference for 28.5°C vs 31.5°C in the presence of CBC. Aversion to 31.5°C stimulated by 10 minute incubation with 5µM CBC is eliminated by knockout of *trpa1b*. Of note, *trpa1b* −/− animals had less baseline aversion to 31.5°C, although this was an inconsistent finding. N = 64 larvae per group. i) Average velocity (mm/s) of larvae in either temperature zone for experiment in panel H. CBC’s specific increase in velocity on the 31.5 side is eliminated with knockout of *trpa1b*. j) Diagram of the *trpv1* mRNA transcript (Ensembl). Our guide RNA (gRNA1) was designed to target exon 5 (red line) k) Sanger sequencing traces from wild-type (above) and *trpv1 −/−* (below) larvae. Yellow highlight indicates overlapping sequence upstream of the mutation. Blue highlight indicates the beginning of overlap downstream of the mutation. The intervening 16bp are lost in the bottom trace. l) Predicted amino acid sequence of wild-type and *trpv1 Δ16* alleles. Red text in the mutant sequence indicates amino acid mismatches occurring after the deletion, and the asterisk marks the new, early stop codon. Mismatches begin at amino acid 243, within the ankyrin repeat domains and before any transmembrane domains of the full protein. m) Colorimetric in-situ hybridization of *trpv1* mRNA in p3 larvae treated with PTU. The stain for this transcript is lost in *trpv1* knockout larva. The inset magnifies the area around the trigeminal ganglion to 200%. Scale bar corresponds to 100µm. n) Effect of *trpv1* homozygous knockout (triangles) on temperature preference for 28.5°C vs 31.5°C in the presence of CBD. Thermal hypersensitivity stimulated by CBD is not affected by knockout of *trpv1*. N = 38-87 larvae per group. o) Average velocity (mm/s) of larvae in either temperature zone for experiment in panel M. CBD’s specific increase in velocity on the 31.5 side is not affected by knockout of *trpv1*. *p < 0.05; **p < 0.01; ***p < 0.001; ns, not significant. Error bars represent SEM. Panels A, C, F, H, N used one-way ANOVA; B, D, G, I, O used two-way ANOVA.

Other cannabinoids have previously been found to activate mammalian TRP channels (Petrocellis et al., 2008). We sought to determine if the pro-nociceptive effects of CBD are conserved in other minor cannabinoids. We found that Cannabichromene (CBC), a close chemical relative of CBD with possible analgesic properties (Sepulveda et al., 2024), also induces thermal hyperalgesia in larval zebrafish at similar doses to CBD (Fig. 2H, I) in a *trpa1b*-dependent manner.

Given that mammalian TRPV1 can be activated by CBD (Bisogno et al., 2001), and is required for thermal hyperalgesia in the mouse (Caterina et al., 1997; Davis et al., 2000), we tested whether zebrafish Trpv1 also contributes to CBD-mediated thermal sensitization. We created a mutant line carrying a 16-bp nonsense mutation in exon 5 of *trpv1* using CRISPR-Cas9, which caused loss of the *trpv1* mRNA (Fig. 2J, K, L). *Trpv1* knockout animals had no difference in CBD-induced thermal sensitization compared to wildtype animals (Fig. 2M, N).

In addition to provoking thermal hyperalgesia, bath application of AITC provokes a Trpa1b-dependent escape response in larval zebrafish which can be measured by an increase in total locomotion (Prober et al., 2008). Given that CBD could provoke similar behavioral effects as AITC in the temperature aversion assay, we tested whether CBD could also mimic AITC’s effects on locomotion by assessing locomotion over a 60-minute period. Individual larvae placed in a single well of a 96 well plate were recorded to measure locomotor activity immediately after adding test solutions (Fig. 3A). Over the first 6 minutes of the experiment, the average speed of vehicle-treated animals was 2.78 ± 0.110 mm/s (Fig. 3B). High concentrations of CBD immediately provoked transient hyperlocomotion increasing average velocity over the first 6 minutes to 3.18 ± 0.088 and 3.25 ± 0.072 mm/s respectively for 4 and 5µM CBD.

**Figure 3).**
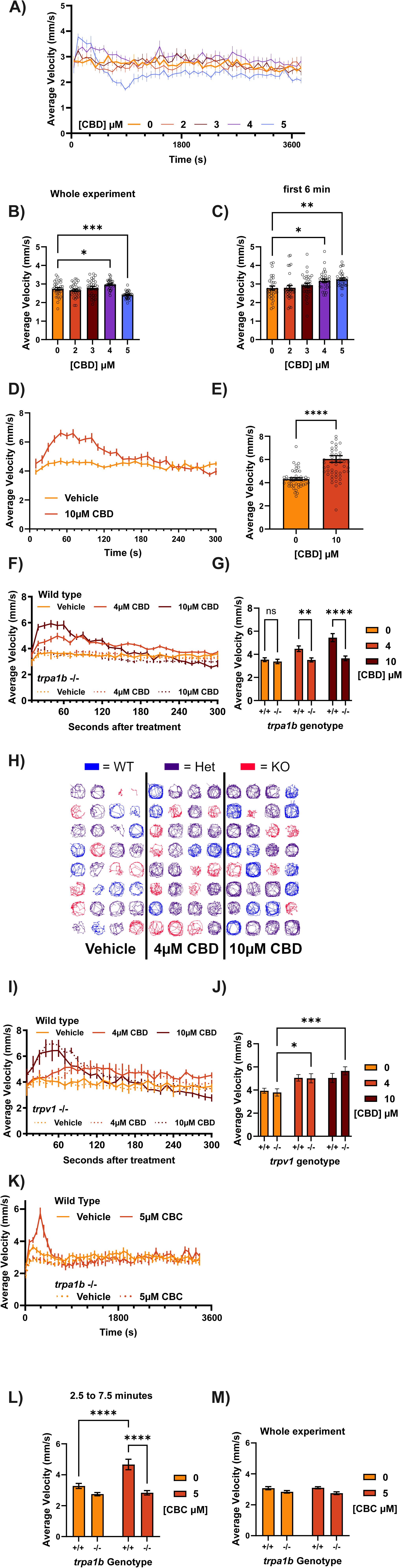
CBD provokes hyperlocomotion dependent on Trpa1b, but not Trpv1. a) Average velocity (10 second binds) of larvae exposed to different concentrations of CBD within the wells of a 96 well plate. Exposure begins less than one minute prior to analysis (time 0). 4 and 5µM CBD induce an acute increase in locomotion, while only 5µM caused a protracted decrease in locomotion afterwards. N = 32 larvae per group. b) Comparison of average velocity (mm/s) of animals over the course of the experiment in panel A (63 minutes). 4µM CBD caused an increase while 5µM caused decrease of the total distance moved by the larvae. c) Comparison of average velocity (mm/s) of animals over the first 6 minutes of the experiment in panel A. Both 4 and 5µM CBD caused acute hyperlocomotion. d) Average velocity (10 second binds) of larvae exposed to vehicle or 10µM CBD within the wells of a 96 well plate maintained at 28.5°C. Exposure begins less than one minute prior to analysis (time 0). 10µM CBD causes acute hyperlocomotion that resolves within 2-3 minutes. N = 48-52 larvae per group. e) Comparison of the average velocity (mm/s) of animals over the first minute of the experiment in panel D. 10µM CBD caused a ∼40% increase (4.31 +- 0.110 to 6.05 +- 0.288 mm/s) in average velocity during this time period compared to vehicle. f) Average velocity (10 second binds) of wild-type (solid lines) or *trpa1b* knockout (dotted lines) larvae exposed to 0, 4, or 10µM CBD within the wells of a 96 well plate. The locomotor responses stimulated by CBD are completely absent in *trpa1b* knockout animals. N = 19-32 larvae per group. g) Comparison of the average velocity (mm/s) of animals over the first minute of the experiment in panel F. 4 and 10µM CBD caused 27% and 54% increases (3.53 +- 0.169 to 4.48 +- 0.245 or 5.45 +- 0.345 mm/s) in the average velocity of wild-type animals compared to vehicle, while *trpa1b* knockout animals lacked any significant increase in average velocity (3.38 +- 0.191, 3.52 +- 0.173, 3.65 +- 0.20 mm/s in vehicle, 4, 10µM CBD groups respectively). N = 19-32 larvae per group. h) Traces indicating the paths taken by individual larvae over the course of the entire experiment in panel F (one of four replicates). Qualitatively, the density of the traces for *trpa1b* knockout animals in the 10µM CBD condition are distinctly lower, corresponding to less total movement. The traces for wild-type and heterozygous animals display greater degrees of thigmotaxis. WT = wild-type, Het = heterozygous for *trpa1b* nonsense allele, KO = heterozygous for *trpa1b* nonsense allele, experiment conducted with offspring from a cross of heterozygous parents. i) Average velocity (10 second binds) of wild-type (solid lines) or *trpv1* −/− (dotted lines) larvae exposed to 0, 4, or 10µM CBD within the wells of a 96 well plate. The locomotor responses stimulated by CBD are unaffected by *trpv1* knockout. j) Comparison of the average velocity (mm/s) of animals over the second minute of the experiment in panel I. 4 and 10µM CBD caused similar increases in the average velocity of wild-type and *trpv1* knockout animals compared to vehicle. N = 19-32 larvae per group. k) Average velocity (10 second binds) of wild-type (solid lines) or *trpa1b* knockout (dotted lines) larvae exposed to 0 or 5µM CBC within the wells of a 96 well plate. CBC produced acute hyperlocomotion lasting over 5 minutes in wild-type, but not *trpa1b* knockout animals, without any protracted decrease in locomotion thereafter. N = 24 larvae per group. l) Comparison of the average velocity (mm/s) of animals over the whole experiment in panel K (56 minutes). 5µM CBC caused no significant changes to average velocity in wild-type or *trpa1b* knockout animals. N = 24 larvae per group. m) Comparison of the average velocity (mm/s) of animals over a five minute period beginning at 2.5 and ending at 7.5 minutes into the experiment in panel K. 5µM CBC caused a 42% increase (3.28 to 4.66 mm/s) in average velocity over this time period in wild-type animals. *Trpa1b* knockout animals had no significant change in average velocity (2.75 to 2.84 mm/s). N = 24 larvae per group. *p < 0.05; **p < 0.01; ***p < 0.001; ns, not significant. Error bars represent SEM. Panels B and C used one-way ANOVA; G, J, L, M used two-way ANOVA; E used a student’s T-test.

After the initial increase, animals treated with 4µM CBD returned to vehicle - treated levels of locomotion for the duration of the assay (Figure 3A). This suggests that after the initial exposure to an algogen, changes in locomotion might not accurately report ongoing nociception, as this dosage of CBD clearly evoked thermal sensitization after a 40-minute incubation where no differences in locomotion between vehicle and CBD (4µM) were observed. However, animals treated with 5µM CBD developed a protracted decrease in locomotion which lasted for the rest of the hour-long recording, such that the average velocity over the entire experiment of 5µM CBD animals was significantly less than vehicle (2.73 ± 0.071 vs 2.40 ± 0.034 mm/s for vehicle and 5µM CBD, Fig 3C). In a separate experiment, animals treated with 10µM CBD had a roughly 40% increase in average velocity (4.31 ± 0.110 to 6.05 ± 0.288 mm/s, Fig. 3D, E) over vehicle during just the first minute of the experiment, which resolved and began to drop below baseline after 5 minutes.

As with thermal sensitization, we found that CBD-induced hyperlocomotion was also Trpa1b dependent (Fig. 3F, G). The average locomotion of wild-type animals over the first minute of drug exposure was increased by 27% and 54% by 4 and 10µM CBD (3.53 ± 0.169, 4.48 ± 0.245, 5.45 ± 0.345 mm/s for 0, 4, 10µM CBD, respectively), but not at all in *trpa1b* −/− animals (3.38 ± 0.191, 3.52 ± 0.173, 3.65 ± 0.200 mm/s for 0, 4, 10µM CBD, respectively). Traces of the path taken by animals in this experiment show that for 10µM CBD, *trpa1b* −/− fish display noticeably less thigmotaxis and overall movement than wildtype or heterozygous siblings (Fig. 3H). CBD-induced hyperlocomotion was not affected by loss of Trpv1 (Fig. 3I, J). The response to CBD differed slightly in terms of the time course, so we chose to compare the second minute of this experiment. In both wild-type and *trpv1* −/− animals, 4 and 10 CBD induced significant increases (3.94 ± 0.216, 5.05 ± 0.278, 5.04 ± 0.395 mm/s wild-type; 3.80 ± 0.313, 5.02 ± 0.395, 5.66 ± 0.355 mm/s *trpv1* −/− for 0, 4, 10µM CBD, respectively) in average locomotion during the second minute of a 5-minute incubation.

Like CBD, 5µM CBC also induced acute hyperlocomotion over the first 5-10 minutes of exposure that was Trpa1b dependent (Fig. 3K, L). However, at this dosage it did not produce a subsequent decrease in locomotion in contrast to CBD (Fig. 3M).

In sum, both CBD and CBC mimic the effects of the Trpa1 agonist AITC in provoking thermal hyperalgesia and hyperlocomotion that is specifically dependent on the noxious signal transducer Trpa1b.

### CBD stimulates activity in *trpa1b*+ sensory neurons by directly activating Trpa1b

If CBD activates Trpa1b in vivo, it should stimulate activity in *trpa1b*+ sensory neurons. To test this, we injected either wild type or homozygous *trpa1b* knockout (KO) zebrafish embryos with pTol2-trpa1b:GCaMP6s, which drives expression of the fluorescent calcium indicator GCaMP6s (Chen et al., 2013) in *trpa1b*-expressing neurons. Fish injected with this plasmid express GCaMP6s specifically in Trigeminal (TG) and Rohon-Beard sensory neurons (Fig. 4A). Consistent with Trpa1b expression, all GCaMP6s-expressing neurons recorded in the trigeminal ganglia responded to AITC with robust calcium influx in WT animals, but not in *trpa1b* −/− animals (Fig. 4B, C, D).

**Figure 4).**
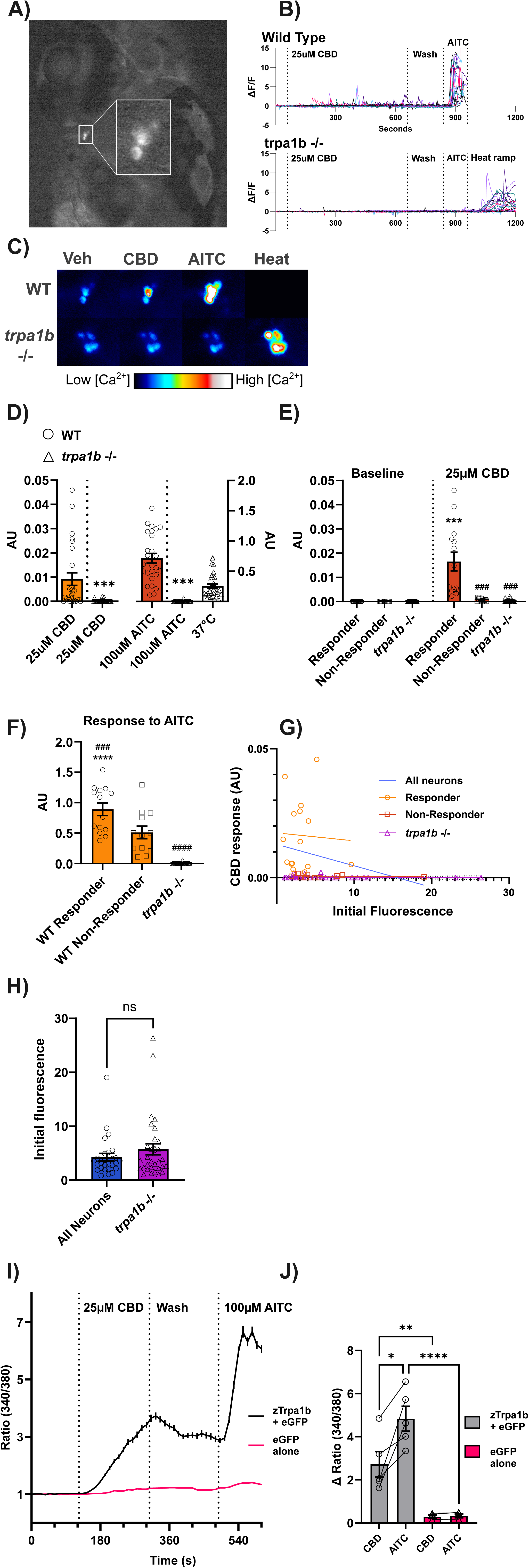
CBD directly activates Trpa1b to activate zebrafish nociceptors. a) Representative P6 larvae used in a calcium imagine experiment, which was injected with pTol2-*trpa1b*:GCaMP6s. Inset magnifies a group of cells expressing GCaMP in a trigeminal ganglion. b) Calcium imaging traces of all recorded neurons (26 AB, 8 fish; 32 *trpa1b* −/−, 9 fish) transiently expressing GCaMP6s. ΔF/F = Change in fluorescence signal relative to the average of the baseline period (average of 28 seconds, 7 frames, prior to the addition of CBD) divided by the average of the baseline. Wild-type neurons display a *trpa1b*-dependent response to 25µM CBD, while *trpa1b* knockout neurons lack responses to CBD and AITC, with a spared response to noxious heat. AITC = Allyl isothiocyanate 100μM. CBD = Cannabidiol. Heat ramp = 24°C to >37°C (ramp duration 60 seconds), held at 37c for 2 minutes. c) Representative fish from wild-type and *trpa1b* knockout groups. Max projection of the response to each chemical stimulus, indicating the peak intensity (calcium concentration) reached by each pixel during a stimulus block. Warmer colors indicate greater calcium concentration. Noxious heat (>37°C) still activates *trpa1b*+ trigeminal neurons after loss of the *trpa1b* protein, although their response to AITC and CBD is lost. d) Quantification of average response to CBD (left Y-axis) and AITC (right Y-axis) in wild-type (circles) and *trpa1b* −/− (triangles) neurons. 14 wild-type neurons had a greater response to CBD than was seen in any of the *trpa1b* −/− cells. No *trpa1b* neurons responded appreciably to AITC, responses to noxious heat were variable but comparable to 100µM AITC. AU = arbitrary unit. Student’s t-test between WT and *trpa1b* −/− groups (excluding 37°C). N=26 wild-type, and 32 *trpa1b* −/−. e) Comparison of response to CBD in 14 ‘responder’ wild-type neurons (circles), 12 non-responder wild-type neurons (squares), and all *trpa1b* −/− neurons (triangles). Average response of responder neurons was significantly greater than baseline (comparison to baseline = asterixes), and from both non-responders and *trpa1b* −/− neurons (comparison to wild-type responders = pound symbols). Neither wild-type non-responder, nor *trap1b* - /- neurons had an average response to CBD that was significantly greater than in the baseline period. N=14 responder wild-type, 12 non-responder wild-type, and 32 *trpa1b* - /-. f) Comparison of response to AITC in 14 ‘responder’ wild-type neurons, non-responder wild-type neurons, and all *trpa1b* −/− neurons. Wild-type CBD responder neurons had a significantly greater average response to 100µM AITC than non-responders (comparison to non-responders = pound symbol) and *trpa1b* −/− neurons (comparison to *trpa1b* −/− = asterixes). *Trpa1b* −/− neurons had a significantly lower response to AITC than wild-type non-responder neurons. N=14 responder wild-type, 12 non-responder wild-type, and 32 *trpa1b* −/−. g) Scatterplot of wild-type neurons, with their initial, background subtracted baseline fluorescence (8 bit intensity, 1-256) on the X-axis and their response to CBD (arbitrary units, see methods) on the Y-axis. There was no relationship between initial fluorescence and the later response to CBD; cells with higher baseline calcium or greater expression of GCaMP did not systematically differ in their responses to CBD. WIld-type neurons overlapped completely with *trpa1b −/−* neurons. Best fit lines created by simple linear regression of the data. Regression line formulas: All neurons, y= - 0.000832x + 0.129, r^2^= 0.0546; Responders, y= −0.000301 + 0.0174, r^2^= 0.00232; Non-responders, y= −1.833e-005x + 0.00070, r^2^= 0.000317; *trpa1b* −/−, y= −1.093e-005x + 0.0002493. h) Comparison of initial fluorescence intensity between wild-type and *trpa1b* −/− neurons. Symbols are individual neurons, data represent the average background-subtracted 8-bit intensity of neurons on the first frame of the baseline period. N= 26 wild-type, and 32 *trpa1b* −/−. i) Ratiometric calcium imaging of HEK-293t cells. Average trace of all HEK 293T cells recorded across 5 coverslips. Coverslips were transfected either with Zebrafish pCDNA3.1-zTrpa1b + pIRES-eGFP (n = 355 cells) or pIRES-eGFP alone (n = 368 cells). 25µM CBD activates zebrafish Trpa1b; cells activated by CBD were also activated by AITC in a plasmid-dependent manner. j) Comparison of the absolute change in 340/380 ratio after application of each solution (CBD vs baseline, AITC vs CBD) for the experiments in panel I. Individual data points are the average for each coverslip. 25µM CBD does not activate zebrafish Trpa1b fully, only generating approximately half the response to 100µM (saturating) AITC. N = 5 coverslips for zebrafish Trpa1b + pIRES-eGFP, 4 coverslips for pIRES-eGFP alone. *p < 0.05; **p < 0.01; ***p < 0.001; ns, not significant. Error bars represent SEM. Panels F and J used one-way ANOVA; E used two-way ANOVA; D and H used a Mann-Whiteney test.

Exposure to 25µM CBD stimulates calcium transients in cell bodies of the wild-type Trpa1b+ TG neurons, although in a much weaker fashion than 100µM AITC (Fig. 4B, C, D, E). This activity begins within five minutes, and peaks between 5 and 10 minutes of a 10 minute application. Unlike for AITC where the response was larger and sustained, responses to CBD were smaller, transient and ‘bursty’. This is potentially due to differences in the potency of these stimuli as well as the uptake kinetics of these compounds in the experimental time frame. The population average response to CBD and AITC was significantly greater in wild type neurons than in *trpa1b* −/− neurons (Fig. 4D), despite the fact that only ∼half (14 of 26) of wild type neurons had a response to CBD which was greater than any seen in the *trpa1b −/−* population. We refer to these cells as ‘responder’ neurons. Responder neurons had a significantly greater response to CBD than to the baseline (vehicle) stimulus, as well as to the CBD response in both non-responder and *trpa1b* −/− neurons. Neither wild-type non-responders, nor *trpa1b* −/− neurons had an average response that was significantly greater than the baseline (Fig. 4E). Neuronal activity stimulated by both AITC and CBD was lost in *trpa1b* −/− animals, while their response to noxious heat (>37°C) was spared, a reflection of the co-expression of Trpa1b with other signal transducers activated by noxious stimuli, such as the heat receptor Trpv1, in nociceptive neurons (Pan et al., 2012; Gau et al., 2013, 2017). These data support a model by which CBD exerts its pro-nociceptive behavioral effects on larval zebrafish by activating *trpa1b*+ nociceptors.

While all wild-type neurons responded to AITC, CBD responder cells had a significantly greater response to AITC than those which did not respond to CBD (Fig. 4F). Given that we cannot control the dose of plasmid each cell receives, it is possible that this difference in AITC response is somehow an artifact of differences in the expression of GCaMP due to plasmid dosing. However, there was no significant relationship between fluorescence intensity on the first frame of the recording and the response to CBD (Fig. 4G) in wild type neurons. The best fit line between the initial fluorescence of wild-type responder neurons and their later response to CBD had an r^2^ of 0.00232, and a slope of -.00031, indicating a poor linear fit and no meaningful correlation between these two variables.

The slope of the best-fit lines for the wild type responders, non-responders, and *trpa1b* −/− neurons were all not significantly different from zero (p = 0.865, 0.686, 0.524 respectively) and none of their slopes were significantly different from each other (p= 0.0556). In addition, the average baseline fluorescence of the *trpa1b* −/− neurons was not significantly different from that of wild-type neurons (Fig.4 H), indicating that there is no systematic difference in activation of the *trpa1b* promoter or in baseline calcium concentration in Trpa1b+ neurons between wildtype and *trpa1b* −/− animals.

When all the wild type cells are considered together, the r^2^ value remains low, such that only approximately 5% of the variance in CBD response might be explained by differences in initial fluorescence. Given that fluorescence will be a function of both GCaMP expression and baseline intracellular calcium, neither factor seems to have strong predictive value for the later response to CBD. We have previously shown (Esancy et al., 2018) that within the larval zebrafish TG, there is heterogeneity in the response of Trpa1-expressing neurons to Trpa1 agonists, where different subsets of Trpa1b-expressing neurons were differentially activated based on stimulus intensity, with some neurons selectively recruited by weak intensity stimuli and others requiring higher intensity stimuli. Our results are consistent with CBD responder neurons representing a sub-population within Trpa1b-expressing sensory neurons with increased sensitivity to Trpa1 agonists.

While these experiments show that sensory neuron responses to CBD are dependent on Trpa1b, they did not allow us to assess whether CBD activates the channel itself, as these neurons could be activated by a downstream effector of CBD. Using calcium imaging again, we measured the change in intracellular calcium in HEK cells transfected with zebrafish Trpa1b in response to 25µM CBD.

Trpa1 is calcium-permeable (Bandell et al., 2004)—opening of the channel will cause a rise in intracellular calcium. HEK-293t cells which were transfected with pCDNA3.0-Trpa1b + pIRES-eGFP (average coverslip fold change of 340/380 ratio = 2.73 ± 0.590 and 4.85 ± 0.578 for CBD and AITC), but not pIRES-eGFP alone (0.291 ± 0.077 and 0.329 ± 0.089 for CBD and AITC), showed robust responses to 25µM CBD and 100µM AITC indicating that zebrafish Trpa1b, consistent with its effects on mammalian TPRA1, is likely to be directly activated by CBD (Fig. 4I, J).

In sum, our data indicate that CBD’s pro-nociceptive behavioral effects, thermal sensitization and hyperlocomotion, are mediated by CBD activating Trpa1b to increase the activity of nociceptive sensory neurons. They also reinforce previous findings that a subset of Trpa1b+ neurons in the trigeminal ganglia are tuned for increased sensitivity to weaker agonists of Trpa1b.

### CBD has anti-nociceptive effects in larval zebrafish

To further explore its analgesic potential, we assayed whether CBD at doses (≤4µM/40min incubation) at or below those required to generate Trpa1b-dependent nocicfensive behavior and/or locomotor depression could inhibit nociception using chemical gradient assays, in which an algogen (AITC (100mM) or acetic acid (1%)) is deposited in a thin layer of agarose on one side of square arena from which it diffuses outward into the surrounding water. The distance from the chemical source each larvae occupies is then measured over time.

Remarkably, forty-minute pre-exposure to 2, 3 or 4µM CBD, which continues during the course of the experiment, dramatically reduced the aversive effects of AITC (Fig. 5B, C) such that by 20 minutes in the assay CBD-exposed fish were not biased away from the AITC source but instead evenly distributed throughout the arena, whereas vehicle treated larvae were strongly biased away from the AITC source. (One sample t-test versus theoretical mean of 50, p = 0.5032, 0.1031, 0.304 for 2, 3, and 4µM CBD respectively). As our locomotor assay showed that none of these doses of CBD reduced baseline locomotion, it is unlikely that the fish were *unable* to avoid AITC due to a locomotor deficiency.

**Figure 5.**
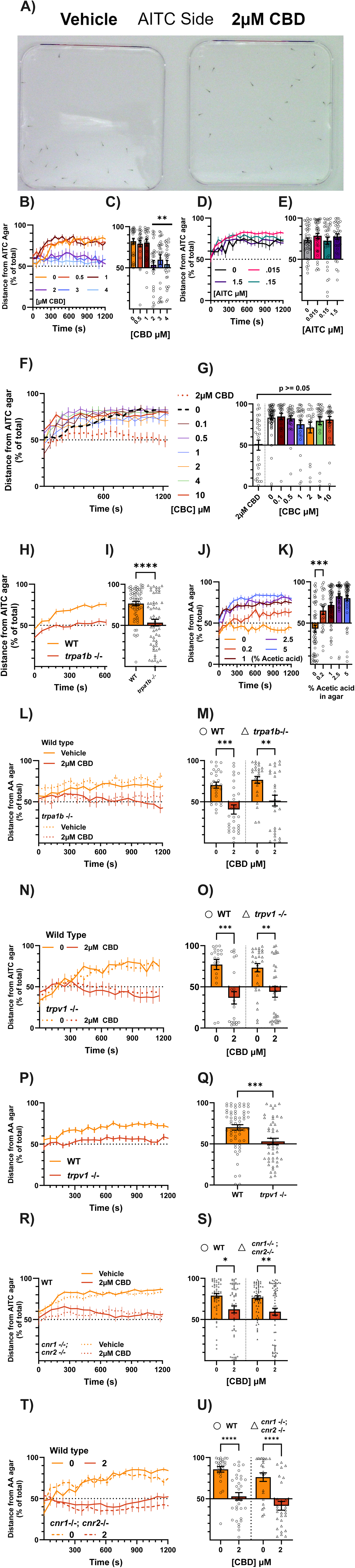
- CBD reduces aversion to chemical irritants. a) Representative image from an AITC aversion experiment with 2µM CBD. The AITC agar lines the top (red marker) of a square petri dish and is allowed to diffuse into solution. At 20 minutes in the assay, vehicle-treated fish avoid the AITC-agar side robustly. Fish treated with 2µM CBD for 40 minutes before being tested in the assay display no net aversion to AITC. b) Dose response curve for 40 minute CBD incubations on aversion to AITC. Data represent the average distance from the AITC side of all (∼32) fish in the square petri dish lined on 3 sides with 0.8% agar E3 medium, and on one side with 100mM AITC in 0.8% agar E3 medium. Distance is normalized to the length of the plate (10cm) where 100% is the maximum distance from the AITC side, and 0% is directly on top of the AITC agar. Every 60th second is plotted. N = 24-32 fish per condition. c) Average distance from AITC agar of each fish in the experiment in panel B at the 20-minute mark (1200 seconds). Data points are individual animals. 2 to 4µM CBD, but not below, prevented aversion to AITC agar. N = 24-32 fish per condition. Additionally, at this time point fish treated with 2 to 4µM CBD, but not below, have an average distance from the AITC agar which is not significantly different from a random distribution (one sample t-test versus theoretical mean of 50%, not shown. P = <0.0001 for 0, 0.5, and 1µM CBD, 0.503 for 2, 0.103 for 3, 0.304 for 4µM CBD). d) Dose response curve for 40 minute AITC incubations on aversion to AITC. Incubation with AITC itself does not reduce aversion to AITC agar. N = 24-32 fish per condition. e) Average distance from AITC agar of each fish in the experiment in panel D at the 20 minute mark (1200 seconds). No significant effects of AITC incubation on AITC aversion at the 20 minute mark. Data points are individual animals. N = 24-32 fish per condition. f) Dose response curve for 40 minute CBC incubations on aversion to AITC. CBC cannot recapitulate the effects of CBD over a wide range of doses. N = 24-32 fish per condition. g) Average distance from AITC agar of each fish in the experiment in panel F at the 20 minute mark (1200 seconds). No significant effects of CBC incubation on AITC aversion at the 20 minute mark. N = 24-32 fish per condition. h) Trpa1b is required for aversion to AITC. Animals treated with vehicle for 40 minutes prior to experiment. N = 64 fish per condition. i) Average distance from AITC agar of each fish in the experiment in panel H at the 20 minute mark (1200 seconds). *Trpa1b* knockout fish (triangles) avoid AITC significantly less than wild-type. Data points are individual animals. N = 64 fish per condition. Additionally, at this time point *trpa1b* −/− fish have a distribution that is not significantly different from chance (one sample t-test versus theoretical mean of 50%, not shown. P = <0.0001 for wild-type, 0.405 for *trpa1b* −/−). j) Cannabinoid receptors Cnr1 and Cnr2 are not required for the effect of CBD upon AITC aversion. Vehicle-treated (yellow) wild-type (solid lines) and double knockout animals for *cnr1* and *cnr2* (dotted lines) avoid AITC, and double knockout animals are fully affected by 2µM CBD (red). N = 64 fish per condition. k) Average distance from AITC agar of each fish in the experiment in panel J at the 20 minute mark (1200 seconds). (TU) *Cnr1 −/−; cnr2 −/−* double knockout animals (triangles) are affected by CBD similarly to wild-type Tubingen animals. Data points are individual animals. N = 59-68 fish per condition. l) Trpv1 is not required for the effect of CBD upon AITC aversion. Vehicle treated (yellow) wild-type (solid lines) and *trpv1 −/−* (dotted lines) avoid AITC, and knockout animals are fully affected by 2µM CBD (red). N = 20-31 fish per condition. m) Average distance from AITC agar of each fish in the experiment in panel I at the 20 minute mark (1200 seconds). *Trpv1* −/− animals (triangles) are affected by CBD similarly to wild-type. Data points are individual animals. N = 20-31 fish per condition. n) Dose response for acetic acid infused in 0.8% agarose EM. Animals pre-treated for 40 minutes with vehicle. Agarose infused with 1% acetic acid or above by volume produces robust place aversion. AA agar = acetic acid agar. N = 25-54 fish per condition. o) Average distance from AITC agar of each fish in the experiment in panel N at the 20 minute mark (1200 seconds). At 20 minutes 0.2% acetic acid agar produces significantly greater aversion than agar without acetic acid. Data points are individual animals. N = 25-54 fish per condition. Additionally, at this time point 0.2% and above acetic acid agar causes greater than chance-levels of aversion, while agar without acetic acid does not (one sample t-test versus theoretical mean of 50%, not shown. P = 0.107, 0.0033, for 0 and 0.2% acetic acid agarose, and <0.0001 for 1, 2.5, and 5% acetic acid agarose, respectively. p) Trpv1 is required for aversion to acetic acid. Animals treated with vehicle for 40 minutes prior to experiment. N = 47-51 fish per condition. q) Average distance from acetic acid agar of each fish in the experiment in panel P at the 20 minute mark (1200 seconds). *Trpv1* −/− fish (triangles) avoid acetic acid significantly less than wild-type. Data points are individual animals. N = 47-51 fish per condition. Additionally, at this time point *trpv1* −/− fish have a distribution that is not significantly different from chance (one sample t-test versus theoretical mean of 50%, not shown. P = <0.0001 for wild-type, 0.443 for *trpv1* −/−). r) Trpa1b is not required for aversion to acetic acid, nor is it required for CBD’s effect on acetic acid aversion. 40 minute pre-treatment with 2µM CBD (red) prevents aversion to 1% acetic acid in both wild-type (solid lines) and *trpa1b* −/− (dotted lines) animals. AA agar = acetic acid agar. N = 26-29 fish per condition. s) Average distance from acetic acid agar of each fish in the experiment in panel R at the 20 minute mark (1200 seconds). *Trpa1b* −/− animals (triangles) are affected by CBD similarly to wild-type. Data points are individual animals. N = 26-29 fish per condition. Additionally, at this time point animals which were pre-treated with 2µM CBD do not significantly avoid acetic acid agar, in contrast to vehicle treated animals (one sample t-test versus theoretical mean of 50%, not shown. P = < 0.0001 for vehicle wild-type and *trpa1b* −/−, 0.105 and 0.826 for 2µM CBD wild-type and *trpa1b* −/− respectively). t) Neither cannabinoid receptor is required for aversion to acetic acid, nor are they required for CBD’s effect on acetic acid aversion. 40 minute pre-treatment with 2µM CBD (red) prevents aversion to 1% acetic acid in both wild-type (solid lines) and *cnr1 −/−; cnr2 −/−* double knockout (dotted lines) animals. AA agar = acetic acid agar. N = 24-33 fish per condition. u) Average distance from acetic acid agar of each fish in the experiment in panel T at the 20 minute mark (1200 seconds). (TU) *cnr1 −/−; cnr2 −/−* double knockout animals (triangles) are affected by CBD similarly to wild-type Tubingen. Data points are individual animals. N = 24-33 fish per condition. *p < 0.05; **p < 0.01; ***p < 0.001; ns, not significant. Error bars represent SEM. Panels C, E, G, K used one-way ANOVA; M, O, S, U used two-way ANOVA; I and Q used a Mann-Whitney test.

In light of our previous results, we questioned whether this antinociceptive effect of CBD—a reduction in sensitivity to a noxious stimulus— was also mediated by Trpa1b. Given that CBD can activate Trpa1b, it may have reduced aversion to AITC by desensitizing the channel or by reducing the net gradient of Trpa1 activators across the plate by simply being present everywhere in solution, thereby giving the fish less reason to avoid any given side of the plate.

To better mimic the intensity of the CBD dosages (2,3 µM) which did not induce thermal sensitization, we chose to incubate animals with AITC beginning with a minimal dose (1.5µM) which consistently causes thermal sensitization and decreased this dose along a log scale. Over a 100-fold concentration range, none of the AITC incubations reduced aversion to AITC agar below control levels (Fig. 5D, E). CBC, which also provoked Trpa1b-dependent behaviors and activates mammalian TRPA1, also failed to replicate the effects of CBD across a wide concentration range (Fig. 5F, G). Therefore, simply activating Trpa1b is not enough to reduce aversion to AITC, and the effects of CBD are specific within its family of phytocannabinoids.

Given that genetic deletion of Trpa1b eliminates aversion to AITC (Fig. H, I), we could not pursue Trpa1b’s role in the effects of CBD by assessing Trpa1b null animals with the AITC aversion assay. Another algogen, acetic acid, produces aversion similarly to AITC, which is also reduced by CBD (Fig. 5J, K). Intriguingly, CBD still reduces aversion to 1% acetic acid agar even in *trpa1b* null mutants (Fig. 5L, M). Collectively, these data suggest that the anti-nociceptive effects of CBD are not mediated by Trpa1b.

Given that TRPV1 has been proposed to mediate some of the analgesic effects of CBD (Costa et al., 2004; Patwardhan et al., 2006), we also tested whether Trpv1 knockout animals lack CBD-induced analgesia. In the AITC aversion assay, *trpv1* −/− animals are still fully affected by CBD (Fig. 5N, O). However, these animals fail to avoid acetic acid (Fig. 5P, Q), precluding us from using the acetic acid aversion assay, but in line with Trpv1’s canonical role in acid-sensing (Leffler et al., 2006; Dhaka et al., 2009; Gau et al., 2013). The cannabinoid receptors Cnr1 and Cnr2, which are conserved in zebrafish, have themselves been implicated in nociception and the analgesic effects of cannabinoids (Agarwal et al., 2007). Larvae lacking both Cnr1 and Cnr2 are fully affected by CBD in both AITC and acetic acid aversion (Fig. 5R, S, T, U).

In sum, doses of CBD which do not induce hyperlocomotion, hypolocomotion, or sensitization to 31.5°C can dramatically reduce aversion to the chemical irritants AITC and acetic acid in a manner which is independent of Trpa1b, TrpV1 and the cannabinoid receptors Cnr1 and Cnr2. Intriguingly, a pronociceptive dose of CBD (4µM) was also able to suppress AITC aversion, suggesting that the pro- and anti-nociceptive effects of CBD are assay dependent.

### Antinociceptive effects of CBD may be masked by its effects on Trpa1

We also assayed the effects of CBD on thermal aversion by measuring place * preference between 28.5°C and the noxious temperature 37.5°C. Neither 40 minute treatment with 1-4µM CBD nor 10 minute treatment with 10µM had any effect on 37.5°C aversion (Fig. 6A, B, C, D), although 40 minutes of 4µM CBD increased average velocity on the 37.5°C side (Fig. 6B) by ∼30% (3.24 ± 0.260 to 4.21 ± 0.474 mm/s) indicative of thermal sensitization and consistent with our earlier findings. We hypothesized that lower dosages of CBD, which did not induce thermal sensitization at 31.5°C, might be able to facilitate nociceptor activity via Trpa1b at 37.5°C, and thus mask the antinociceptive effects of CBD. This would not necessarily be reflected with a change in behavior due to a ceiling effect, as the fish already robustly avoid 37.5°C. If this were the case, then antinociceptive properties of CBD may emerge in *trpa1b* null animals. Therefore, we tested whether the highest effective antinociceptive dose of CBD (3µM) could inhibit thermal aversion in Trpa1b null animals. Remarkably, CBD (3µM) which had no significant effect on 37.5°C aversion in WT animals (83.6% ± 2.07, 73.8% ± 3.74 for 0 and 3µM CBD respectively, Fig. 6E), completely inhibited 37.5°C aversion in *trpa1b* null animals (88.9% ± 1.82, 51.2% ± 5.32 for 0 and 3µM CBD respectively). Importantly, there was no difference in 37.5°C aversion between vehicle-treated WT and *trpa1b* null animals. This data suggests that the pro-nociceptive effects of CBD, mediated by Trpa1b, can antagonize and mask its anti-nociceptive properties.

**Figure 6.**
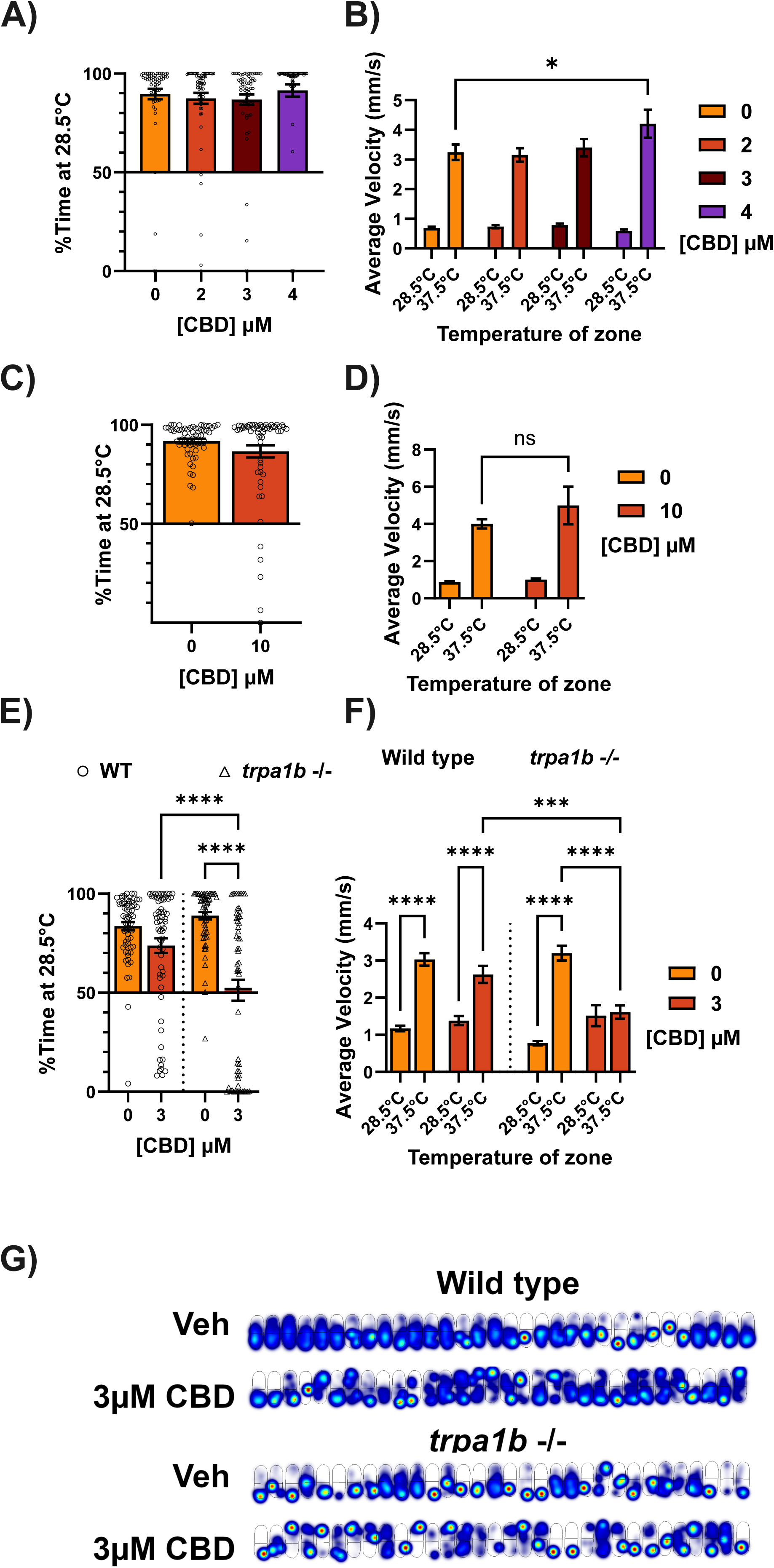
- Trpa1 agonism masks an analgesic effect of CBD. A) Dose response for 40 minute incubation of CBD prior to testing temperature preference for 28.5°C vs 37.5°C. Data represent average time spent on the 28.5°C side across all fish as a percentage of total trial duration (4 minutes). 37.5°C is highly aversive after vehicle and 2-4µM CBD treatments. N = 64 larvae per group. B) Average velocity (mm/s) of larvae in either temperature zone for experiment in panel A. Animals have much higher average velocities on the 37.5°C side, and 4µM CBD exacerbates this effect. C) 10 minute pre-treatment with 10µM CBD does not affect aversion to 37.5°C. Data represent average time spent on the 28.5°C side across all fish as a percentage of total trial duration (4 minutes). N = 64 larvae per group. D) Average velocity (mm/s) of larvae in either temperature zone for experiment in panel C. Animals have much higher average velocities on the 37.5°C side, which is unaffected by 10 minute treatment with 10µM CBD. E) Trpa1b agonism by CBD masks a separate analgesic effect on temperature preference which is not mediated by Trpa1b. Data represent average time spent on the 28.5°C side across all fish as a percentage of total trial duration (4 minutes). *Trpa1b −/−* (triangles), but not wild-type animals, lose aversion to 37.5°C after 40 minute pre-treatment with 3µM CBD. N = 64 larvae per group. Additionally, only *trpa1b* −/− fish treated with 3µM CBD have a distribution that is not significantly different from chance (one sample t-test versus theoretical mean of 50%, not shown. P = <0.0001 for vehicle treatments, wild-type 3µM CBD, 0.8203 for *trpa1b* −/− 3µM CBD). F) Average velocity (mm/s) of larvae in either temperature zone for experiment in panel E. Wild-type and *trpa1b* −/− animals have greater velocity on the 37.5°C side after vehicle treatment, but only *trpa1b −/−* animals have equal velocities on both sides after 40 minute treatment with 3µM CBD, consistent with a lack of escape response to 37.5°C. G) Heat maps indicating the cumulative position of the animals over the course of the experiment in panel F. Brighter/hotter colors indicate more time spent in those pixels. The 28.5°C zone is below the dark line across the center of the arena, and the 37.5°C zone is above this line. Wild-type animals exhibit a bias towards the 28.5°C side with vehicle and 3µM CBD treatment, while *trpa1b* −/− animals treated with 3µM CBD are spread across both temperature zones. *p < 0.05; **p < 0.01; ***p < 0.001; ns, not significant. Error bars represent SEM. Panel A used one-way ANOVA; B, D, E, F used two-way ANOVA; C used a Mann-Whitney test

## Discussion

In this study, we have demonstrated that CBD has both pro- and anti-nociceptive behavioral effects—depending on concentration and nociception test—in larval zebrafish. At relatively high doses, CBD mimics the effects of the Trpa1 agonist AITC on temperature sensitivity and locomotion, but at lower doses can block aversion to chemical algogens. Intriguingly, an analgesic dosage of CBD was able to reverse heat aversion in Trpa1b null animals. Our findings indicate that the pro-nociceptive, but not the anti-nociceptive, effects of CBD are mediated by Trpa1b, and that agonism at Trpa1b may counteract CBD’s analgesic effects. Additionally, putative targets for CBD-mediated antinociception such as Trpv1 or the cannabinoid receptors Cnr1 and Cnr2 had no apparent role in CBD’s effects on pain sensitivity.

In our study, CBD’s pro- and anti-nociceptive effects occurred at different dose ranges. Sensitization to 31.5°C began only at or above 4µM for 40-minute incubations, and transient hyperlocomotion also only occurred at or above 4µM, with higher doses increasing these effects. Interestingly, at this dosage thermal sensitization occurred on a time scale long after the hyperlocomotive effects of CBD have resolved. While both thermal sensitization and hyperlocomotion were dependent on Trpa1b, activation of Trpa1b by CBD at this dosage was not sufficient to induce immediate thermal sensitization.

Meanwhile, lower concentrations of CBD (2-3µM), which did not induce sensitization to 31.5°C or locomotor effects, were completely effective in blocking AITC and acetic acid aversion. Therefore, in a brief dose window above 1µM and below 4µM for 40 minutes—whose timing is consistent with CBD’s reported peak concentration after bath application (Achenbach et al., 2018)—CBD produces maximum analgesic effects for algogen aversion without evoking Trpa1b-dependent nociception. Further, 3µM CBD, which was ineffective at reducing aversion to 37.5°C in wild type animals completely abolished aversion in *trpa1b* null larvae. The picture that emerges is one of two closely overlapping dose response curves, one for CBD’s pro- and one for its anti-nociceptive effects, whose relative strength at a given dose are dependent on the modality and intensity of the pain stimulus. Illustrative of this point is the fact that 4µM, which was pro-algesic in the thermal aversion and locomotion assays, was antinociceptive in the AITC aversion assay. Our evidence suggests that Trpa1 agonism plays a modulatory (sensitizing) role in thermal nociception; Trpa1b was not required for aversion to the acutely noxious temperature 37.5°C, but it was required for sensitization to a milder temperature, 31.5°C. On the other hand, prior Trpa1 agonism did not affect aversion to AITC. Trpa1’s divergent sensitizing potential in these two contexts—thermal and chemical nociception—may underly CBD’s ability to display both pro and anti-nociceptive effects at the same dose in two separate assays of pain sensitivity. We were unable to identify a molecular target necessary for CBD’s antinociceptive effects. A putative mechanism for cannabinoid-mediated analgesia involves the cannabinoid receptors Cnr1 & 2 (Finn et al., 2021). Both are Gi/Go_α_ -coupled GPCRs, although their expression patterns are dissimilar (Stella, 2023). Cnr1 is canonically expressed in neurons of the central and peripheral nervous system, where it has been implicated in pain sensation by acting as a negative regulator of activity in sensory neurons. Cnr2 is canonically expressed in cells of the immune system—although an increasing body of evidence suggests its expression in the nervous system and its involvement in brain function, particularly under conditions of pathology (Guerrero-Alba et al., 2019). Both genes are found in the larval zebrafish with similar expression patterns as in mammals (Son and Ali, 2022).

CBD has low affinity for CB1 and CB2 receptors but has been proposed to act as a low-potency negative allosteric modulator of CB1 (Stella, 2023). In our study, neither cannabinoid receptor was required for any of CBD’s behavioral effects, whether pro- or anti-nociceptive. These results are in line with studies which found no effect of CB1 or CB2 receptor antagonists on CBD-mediated analgesia in the mouse (Silva-Cardoso and Leite-Panissi, 2023).

Another putative mechanism for cannabinoid-mediated analgesia involves TRP channels, and particularly desensitization of TRP channel-expressing sensory neurons (Muller et al., 2019). Agonists of TRP channels can induce desensitization by modification of channel function, as well as by internalization and physical removal of the channel from the plasma membrane (Gordon-Shaag et al., 2008). Indeed, CBD has been shown to desensitize the response of rat TRPA1 and TRPV1 to later stimulation by their agonists (De Petrocellis et al., 2011, Akopian et al., 2008). Given TRP channels are often polymodal and may integrate multiple sensory modalities, desensitization stimulated by just one modality could lead to a reduced response to other nociceptive stimuli. Therefore, activators of nociceptors could induce prolonged hyposensitivity— and presumably reduced pain. In line with this theory, Capsaicin and similar compounds which activate pain neurons are used in clinical settings to control pain (Fattori et al., 2016).

In our study, both the TRPA1 agonists CBC and AITC failed to replicate the effects of CBD in the AITC aversion assay. Furthermore, Trpa1b was not required for CBD mediated inhibition of acetic acid aversion. Given that CBD was only effective in reducing aversion to 37.5°C after Trpa1b was eliminated, and our data suggests that Trpv1 was not involved in either CBD pro- or anti-nociception, our data do not support a model whereby CBD-mediated antinociception involves activation of, or desensitization of, these TRP channels, but instead likely relies on another target(s). However, this does not rule out that over a longer time scale than assessed in this study, CBD could desensitize TRP channels and thus enhance its anti-nociceptive properties. Similar to our finding that *trpa1b* knockout reversed CBD thermal sensitization and revealed an antinociceptive effect of CBD on noxious heat (37.5°C) aversion, a previous study found that treatment with an antagonist of Trpa1 could attenuate CBD-mediated thermal hyperalgesia, and accentuate CBD-mediated analgesia, in a rat model of osteoarthritis (Mlost et al., 2021). This study found lower doses of CBD to be analgesic while higher doses were pro-nociceptive, similar to ours. This could be indicative of a conservation of mechanism between zebrafish and higher vertebrates where the pro-nociceptive and anti-nociceptive properties of CBD dose-dependently antagonize each other. Furthermore, temperature appears to be a major determinant in the nociceptive potential of CBD. A dose of CBD (3µM) that was not pro-nociceptive at room temperature (∼27°C) and was anti-nociceptive in our algogen aversion assays became pro-nociceptive in wild type larvae, as opposed to *trpa1b* KO, at 37.5°C. As the pro-nociceptive effects of CBD are dependent on Trpa1b, these findings align with studies in mice showing a role for TRPA1 in heat nociception (Hoffmann et al., 2013; Nozadze et al., 2016; Vandewauw et al., 2018).

As to what mechanisms may mediate CBD-mediated analgesia, there are number of potential targets that can be explored in future studies. These include GPR55, an orphan GPCR, that has been considered to be a third cannabinoid receptor, as its activity is modulated by anandamide (Lauckner et al., 2008) and other cannabinoids. It has also become implicated in CBD’s ability to combat seizures (Rosenberg et al., 2023) as well as in pain pathways independently of CBD (Singh et al., 2024). Other potential targets include the serotonin receptor 5HT_1A_, the proliferator-activated receptor alpha (PPAR-α), opioid receptors, orphan GPCR’s such as GPR18 and GPR119, and other TRP channels (Mlost et al., 2020; Britch et al., 2021; Stella, 2023). Furthermore, CBD’s polypharmacology opens the possibility for combinatorial effects or interactions between multiple targets which are greater than the sum of their parts.

Both animal and human studies have found inconsistent efficacy from CBD as an analgesic (Silva-Cardoso and Leite-Panissi, 2023). In this study, CBD was effective at reducing aversion to chemical algogens in wild-type animals, while it was ineffective at reducing aversion to elevated temperatures. These results are consistent with the modality of pain being an important factor in the efficacy of CBD as an analgesic. Our findings suggest that studies should ideally test CBD across a range of doses and independent pain modalities to form the most accurate picture of how CBD modulates nociception, as our data suggests a dose-dependent antagonism between the pro- and anti-nociceptive properties of this compound. Furthermore, route of administration should be considered. While it is not possible to administer CBD systemically via immersion in mammals as in larval zebrafish, it is likely that the relative concentration of CBD for instance in the peripheral versus the central nervous system could play a role in the effectiveness of CBD as an analgesic. Indeed, a recent study utilizing a novel, long-term ad-libitum oral administration protocol for CBD consumption in mice demonstrated significant improvement in pain phenotypes from CBD gelatin (Abraham et al., 2020).

Cumulatively, our results demonstrate that CBD activates zebrafish Trpa1b to cause its pro-nociceptive behavioral effects. Meanwhile, non Trpa1b-dependent mechanisms underlie a separate anti-nociceptive effect of CBD. Further studies may investigate the effects of CBD and other cannabinoids on pain sensation in the absence of Trpa1 to dissect away parallel, countervailing effects of this promiscuous molecule.

## Acknowledgements

We thank Dr Wolfram Goessling for his generous gift of the Cnr1 and Cnr2 knockout lines.

## Contributions

A.D. and Ben L. conceived the project. A.D. and Bryce L. designed research. Bryce L, Q.B, S.R, G.S, K.E performed research. Bryce L analyzed data. Bryce L, A.D and Ben L wrote the manuscript.

